# Gain and Loss of Acquired Doxycycline Resistance in *Lactiplantibacillus plantarum*: An Adaptive Laboratory Evolution Study

**DOI:** 10.64898/2026.03.17.712413

**Authors:** Bryce Pierce, Olivia Love Harrison, Brooks Floyd, Agnes Moriarty, Kwadwo Antwi-Fordjour, Robert A. Hataway, Brad C. Bennett

## Abstract

Probiotics interact in a mutualistic way with humans. Exposure of probiotics to sublethal concentrations of commonly prescribed antibiotics can cause resistance to arise. Over 1000 generations, we performed an adaptive laboratory evolution (ALE) experiment to determine if the fitness of the probiotic *Lactobacillus plantarum* is altered by exposure to doxycycline (DOX) at 1/10 MIC. Compared to the original culture, *L. plantarum* exposed to sublethal DOX acquired modest resistance (∼4-fold) over time. When the selection pressure was removed, resistance was lost rapidly, in ∼50 generations. This suggests that resistance, once acquired, is not fixed. The mechanism by which resistance is acquired and subsequently lost was investigated by whole genome sequencing (WGS). Analysis of the single nucleotide variants (SNVs) identified in the WGS of the generation 1000 DOX-treated cultures reveal 16 distinct variants across 15 genes. Two of these variants are in the *rpsJ* gene, which encodes ribosomal protein s10, a component of the 30S ribosomal subunit, and result in non-synonymous mutations (H56Y and S94N). This gene has been previously reported to harbor mutations associated with tetracycline class resistance, including DOX and tigecycline. WGS of archived cells from generations 350 and 750 reveals that one of these *rpsJ* variants (H56Y) arises early in the experiment. Additional *rpsJ* variants at position 57 (K57M, K57I) could be identified within intermediate generations. Most *rpsJ* SNVs identified from the WGS could be verified by colony PCR and Sanger sequencing. Importantly, no *rpsJ* variants are observed in the original culture sequences or in same-generation controls.

**Importance:** Probiotics are beneficial bacteria often found in fermented foods and beverages. When consumed, they can colonize the gut and provide several advantages. Antibiotics, even at levels too low to kill bacteria, can exert pressure that leads to antibiotic resistance. Though well-studied in harmful bacteria, it is less understood what effects this may have on probiotics. Using experimental evolution, we grew a popular probiotic, *Lactiplantibacillus plantarum*, in the presence of a low concentration of doxycycline, one of the most highly prescribed antibiotics in the US. Over the course of 5 months, roughly a thousand generations, the probiotic became ∼4-fold more resistant; once we removed the antibiotic and continued the experiment, this resistance disappeared rapidly. DNA sequencing identified variants in the *rpsJ* gene, which encodes a protein important for translation. As doxycycline interferes with this process, the *rpsJ* variants may underlie the emergence of modest resistance we observe.

---

Probiotics are live microbes in food, beverages, or supplements that, when administered or consumed in adequate amounts, can provide a health benefit to the host (1). They colonize the animal intestinal tract and become symbiotic members of the gut microbiota. Beneficial functions range from assisting in the fermentation of indigestible carbohydrates, the secretion of useful metabolites such as short chain fatty acids (SCFAs), and the prevention of infection by pathogens due to barrier protection (1, 2). Probiotics are often used clinically as an adjuvant therapy for gastrointestinal maladies such as irritable bowel syndrome and antibiotic-associated diarrhea (3). There is recent immense interest in how certain probiotics can influence the gut-brain axis, a bidirectional pathway bridging the gut microbiome with neurological function (4). Evidence of benefits of these “psychobiotics” have been in the treatment of neurodegenerative diseases, such as Alzheimer’s Disease (4), and psychiatric disorders, for example, Major Depressive Disorder (5). There are dozens of species of probiotics, each represented by numerous strains. They mainly fall into 3 categories: the genera *Bacillus*, *Lactobacillus,* or *Lactiplantibacillus* (6); the genus *Bifidobacterium*; and other species from *Enterococcus faecalis* to the yeast *Saccharomyces boulardii* (2). *Lactobacillus* is a large genus of Gram positive, non-spore forming bacilli that produce lactic acid and thus belong to the Lactic Acid Bacteria (LAB) supergroup (2). The *Lactobacillus* genus is divided into numerous groups based on fermentative capabilities and/or 16s *rRNA* sequences (7, 8). A recent reorganization and reclassification propose splitting *Lactobacillus* into 25 total genera. Specifically, the plantarum group is now reclassified under the new genus name *Lactiplantibacillus*, with about 15 distinct species (6). *Lactiplantibacillus plantarum* is one of the most well-studied LAB species and can be naturally found in dairy, meat, and vegetable products undergoing fermentation, such as yogurts, aged sausages, and kimchi (8). *L. plantarum* has a high tolerance for acidic environments, and, as such it can survive passage into the gut and colonize there as a member of the gut microbiota. Many positive health benefits have been attributed to *L. plantarum*, a significant one being in the treatment of inflammatory bowel disease (IBD), where it may be able to suppress inflammation and regulate the immune response (3).

Antibiotics are critically important therapeutics in the prevention, treatment, and eradication of many infectious diseases caused by pathogenic bacteria. Their discovery and subsequent development of antimicrobials based on them had arguably the most significant impact on human health outcomes in the 20^th^ century; indeed, the discovery and eventual large-scale manufacturing and clinical deployment of penicillin alone is attributed to saving millions of lives (9). Microbes can evolve resistance to antibiotics, known as antimicrobial resistance or AMR, using a myriad of mechanisms and can do so rapidly. In fact, the first documented case of penicillin resistance was reported in 1945, only 4 years after its clinical use became widespread (10). This suggests that the “tools” required for these resistance mechanisms (resistance genes) were genomically already in place and ancient, easily mobilized on plasmids and transposons (11, 12). The emergence of AMR in just 6 common human microbial pathogens, such as the methicillin-resistant strain of *Staphylococcus aureus* (MRSA), causes >35,000 deaths and cost >4 billion USD annually (13). A recent report projects the annual global health burden caused by AMR pathogens to be ∼1.5 million deaths, with the economic burden to be ∼66 billion USD (14).

Many sources have been attributed to the precipitous rise in AMR. However, a consensus is that the over-prescription, overuse, and misuse of antibiotics in human health settings is at least partially to blame. Most antibiotics prescribed for human use are done so in outpatient settings (15), and nearly 30% of those have been described as unnecessary (16). Whie antibiotic consumption and consumption rates have been fairly flat or rising modestly in high-income countries, these metrics rose dramatically in low to middle income nations from 2000-2015, especially for broad-spectrum penicillins (17). Tetracyclines, and specifically doxycycline, is the 4^th^ most prescribed antibiotic in the US (15). Tetracycline is a broad spectrum antibiotic, with activity against a large range of Gram positive and Gram negative bacteria. Its mechanism of action is to stall protein synthesis by binding the 16s subunit of the 30S ribosome near the A site, preventing aminoacyl-tRNAs from accessing the site (18), (19). Tetracycline was discovered in 1948, however, the first report of a clinically resistant microbe, a strain of *Shigella*, was isolated in 1953 (20). Semisynthetic approaches resulted in modified tetracyclines, including a 6-deoxy, 5-hydroxytetracycline (doxycycline; DOX) that was released in 1967 and marketed under the name vibramycin (10, 20). However, reports of DOX resistance occur as early as 1976 (21). Tetracycline class resistance genes mainly fall into two categories: those encoding proteins associated with or are membrane protein pumps (such as *tet(L)* and *tcr3*) and are responsible for mediating the efflux of antibiotic to outside the cell; and, those that encode cytoplasmic proteins that protect the ribosome from antibiotic binding (22). These ribosomal protection proteins (RPPs) in prokaryotes, the most well-studied being TetM, are related to elongation factors in eukaryotes and essentially block a tetracycline antibiotic from binding the A site or can even dislodge one previously bound(23).

*L. plantarum* is generally susceptible to β-lactam antibiotics, such as penicillin, however, there have been reports of β-lactamase (*bla*) genes in some strains (22). In contrast, a recent study revealed that nearly 80% of tested Lactobacilli strains are inherently resistant to vancomycin (24). Indeed, resistance genes for almost every major antibiotic class, with some strain variability, have been identified across the Lactobacilli. This is certainly the case for tetracycline antibiotics, with *tet* genes found in strains of many species within the Lactobacilli (22, 25). In 2019, a genus-wide study of antibiotic susceptibility, ARGs, and ARG phenotype-to-genotype relationships across 182 type strains of Lactobacilli was reported. Exactly half of the strains had resistance to tetracycline, with 7 of the 10 *L. plantarum* phylogroup strains tested being resistant(24). Genome sequences were available for 161 of the 182 type strains, and these were aligned against ARG resistance genes in the Comprehensive Antibiotic Resistance Database (CARD) (26). Eighteen tetracycline resistance genes were identified across the 161 type strains, highlighted by the presence of RPP genes (such as *tet(M)*) and/or efflux pumps (such as *tet(L)*) in most of the resistant strains. Surprisingly, in the *L. plantarum* phylogroup, none of the 6 major RPP or efflux genes (*tet(M)*, *tet(S), tet(Q), tet(W), tet(L),* and *tet(P)*) were identified in any of the strains (24). It could be that one of the other 12 tetracycline resistant genes cross-referenced by alignment in CARD or lower frequency genes, such as *tet(K)* and *tet(O)* (25, 27), were present. Another possibility are resistance genes that fall outside of the *tet* or *tcr* families. Multiple studies examining bacterial resistance to tigecycline, a glycylcycline derived from tetracycline that binds with higher affinity to the 30S ribosomal subunit(28), have found that mutations in certain efflux pumps (such as AcrAB-TolC) or in 16s rRNA and/or ribosomal proteins are most common (29). For the latter, these include mutations in the *rpsC* and *rpsJ* genes, which encode the S3 and S10 ribosomal proteins, respectively. Although these proteins do not contact the ribosomal A site directly, they do interact with helices h31 and 36 of the 16s rRNA. All the S10 mutations reported reside within a long, unstructured loop (residues 50-60 using *E. coli* numbering), which contacts the h31 helix, and it may be that mutations here influence packing of h31 near the A site, weakening tigecycline binding there. These tigecycline resistance studies include both Gram positive and Gram negative species, including *Staphylococcus aureus*, *Streptococcus pneumoniae*, *Escherichia coli*, and several others (29–34). Mutations in *rpsJ* linked to DOX resistance has also been reported in a few species, such as *Staphylococcus aureus*, *Neisseria gonorrhoeae*, and *Vibrio cholerae*, mostly from adaptive laboratory evolution (ALE) experiments where cultures are passaged in the presence of sublethal concentrations of DOX for many generations (21, 30, 35–37).

To our knowledge, there hasn’t been a published study of an ALE experiment for *L. plantarum* in the presence of a tetracycline class antibiotic. Previous ALEs with other antibiotics, such as streptomycin (38, 39), gentamicin (40, 41), and ampicillin(42), have been reported (**Table 1**). Here we present the results of an ALE experiment with *L. plantarum* where it was continually cultured in the presence of sublethal DOX for ∼1000 generations. Modest resistance was acquired, and sequencing revealed that several *rpsJ* variants arose over the course of the experiment, all resulting in non-synonymous mutations. These variants could not be identified in sequences of either the original cultures or same-generation controls. DOX resistance was rapidly lost when the antibiotic was removed after generation 1000. Thus, *rpsJ*-mediated resistance is reversible, suggesting certain mutations of this gene have a high fitness cost.

**Table 1.** Previous ALE Studies with *L. plantarum* strains.

| Strain | Antibiotic | Concentration | Duration (in days) | Fold Increase in Resistance | Reference |
| --- | --- | --- | --- | --- | --- |
| P-8 | Ampicillin | 0.5 MIC | 365 | 32x | Cao et al (2020)(57) |
| ATCC 14917 | Gentamicin | IC<sub>50</sub>* | 30 | 128x | Dong et al (2019)(41) |
| MCC 301 | Gentamicin | 0.25 MIC* | 30 | 16x | George et al (2017)(40) |
| ATCC 14917 | Streptomycin | IC<sub>50</sub>* | 25 | 8192x | Zhang et al (2018)(39) |
| IMAU 60045 | Streptomycin | IC<sub>50</sub>* | 30 | 1024x | Shao et al (2015)(38) |

\*Antibiotic concentration in the ALE was adjusted every 5 days based on new MIC or IC<sub>50</sub> determination.

## Materials and Methods

### Bacterial strain, growth media, and reagents

We acquired a lyophilized stock of *Lactiplantibacillus plantarum* subspecies *plantarum* (Orla-Jensen) from the American Type Culture Collection (ATCC), with the ATCC strain designation 14917, known hereafter as *Lp14917*. The cells were reconstituted and subsequently cultured in Mann-deRosa-Sharpe (MRS) *Lactobacillus* media (Hardy Diagnostics; Santa Maria, CA). Cells were regularly frozen (“archived”) by mixing an aliquot of an overnight culture with 60% (*v/v*) sterilized glycerol, for a final concentration of 15% (*v/v*) of cryoprotectant. Cryoprotected cells were then placed in a −80 freezer for long term storage. Doxycycline (Thermo Scientific; Waltham, MA) was purchased as a powder and reformulated in dH_2_O.

### Establishing DOX susceptibility and estimating a MIC for *Lp14917*

We used disk diffusion assays to determine the susceptibility of *Lp14917* to DOX. Different concentrations of DOX were prepared by mixing antibiotic powder with the proper volume of dH_2_O. In triplicate we inoculated MRS agar plates with an overnight culture of *L. plantarum* and subsequently placed DOX-soaked discs to the central region of three quadrants of the plate with the fourth containing a disc soaked in 1 x TBS as a control. The DOX-soaked discs varied in concentration per quadrant and thus inhibitory strength. Plates were then incubated for 48 hours at 35°C. After incubation, we observed halos of no growth around the discs, or zones of inhibition (ZOIs), for some of the concentrations tested. Measurable ZOIs were only observed in quadrants with disks soaked in DOX at > 30 µg/mL. This was interpreted as a rough estimate of the susceptibility of this strain to DOX and guided our subsequent broth dilution assays to determine a MIC value.

We initially investigated this using the macrobroth dilution method, with a dilution series that covered ∼100-fold range: 825, 413, 206, 103, 52, 26, 13, 6.5, 3.2, 1.6, and 0.8 µg/mL. We inoculated 5 mL cultures of MRS broth containing known concentrations of DOX with a 5% inoculum of an overnight culture of *Lp14917*. We then incubated in a shaking incubator at 225 rpm at 35°C for 24 hours. To determine total growth and changes in growth rate, at the 24-hour mark, we pipetted 100 µL of culture into wells of a 96 well plate and recorded optical density measurements at 600nm (OD_600_) in an Infinite M200 Pro microplate reader spectrophotometer (Tecan US; Charlotte, NC). This provided a general range for the MIC and informed the setup for our microbroth dilution assays so we could determine the exact MIC as well as an inhibition profile.

For the microbroth dilution assays, we prepared 96 well plates with 100 µL of MRS broth and serially diluted concentrations of DOX in each well, using the same dilution series as above. We then inoculated each well with 100 µL of a 5% inoculum of an overnight culture of *Lp14917* and allowed to incubate at 35°C for 24 hours. OD_600_ measurements were recorded, and total growth and growth rate was compared to control wells. We used both visual inspection and OD_600_ values to assess growth after 24 hours of incubation (43). We roughly determined the MIC by observing apparent visual growth in wells at the highest antibiotic concentration. Visual inspection alone indicated a MIC of 26 µg/mL. Next, for all wells, we compared the 24-hour OD_600_ values to the time 0 value to determine a growth rate. As visual inspection is the standard for establishing MICs (43), we estimated the MIC at 30 µg/mL, though the real value is likely a range between 26 – 52 µg/mL. Importantly, this is well within the range that has been previously reported, where an average MIC for DOX and *L. plantarum* (representing several different strains) was found to be 26.8 µg/mL (24). Thus, our working MIC to guide the ALE experiment (30 µg/mL) appears to be historically accurate.

### Adaptive laboratory evolution experiment

A small aliquot of frozen stock of *Lp14917* was inoculated onto an MRS agar plate, and the plate was incubated at 35°C for 48 hours. Three spatially separated colonies were chosen and inoculated into 3 separate tubes of 3 mL of MRS broth. These were allowed to incubate overnight, and we then used these cultures as our generation zero (Gen0) cultures and starting point for the ALE experiment. Aliquots of these colonies were frozen as our Gen0 cells for the archive.

Each of the 3 cultures were used to inoculate 2 tubes with 3 mL of MRS broth, one designated control and one designated DOX, the culture exposed to a sub-lethal concentration of doxycycline. A 5% *v/v* inoculum was used. Prior to inoculation, doxycycline was added from a 1000X stock to the DOX tubes, bringing the final concentration of DOX in these tubes to 3 µg/mL, which is 1/10^th^ of the MIC. The control tubes received no antibiotic. The 6 culture tubes (3 controls + 3 DOX-exposed) were incubated 35°C with shaking (225 rpm) for no more than 24 hrs. The following day, the inoculation procedure was repeated, with a previous day’s culture used to inoculate the current day’s media, and the cultures placed back into the shaking incubator for overnight incubation. This process of continual culturing with daily propagation went on for 5 months, or ∼1000 generations. The experiment was continued as normal for another 375 generations except that no doxycycline was included in the MRS broth for the DOX-exposed cultures.

### Fitness test assays

To determine if DOX-exposed cultures gained resistance to DOX during the ALE, biweekly fitness tests (∼100 generations) were performed in 96 well plates (**Figure 1**). Each row on the plate corresponded to one bacterial colony (3 controls + 3 DOX-exposed), and each column represented a concentration of DOX (using the above concentration series). The concentration of DOX in each column decreases by a factor of ½ as one moves from the leftmost to the penultimate right column. The rightmost column is a positive control for growth and contains no DOX. To each well, 100 µL of MRS broth was added. A stock concentration of DOX was prepared in deionized water at 1.65 mg/mL. Subsequently, 100 µL of DOX stock was added to the first column of the plate, mixed, and then serially diluted across the plate until the 11^th^ column, with the highest DOX concentration in column 1 at 825 µg/mL and the lowest in column 11, at 0.8 µg/mL. The 12^th^ well is untreated, with only the MRS broth. Each well is then inoculated with 5 µL (5% *v/v*) of the ALE cultures.

**Figure 1.**
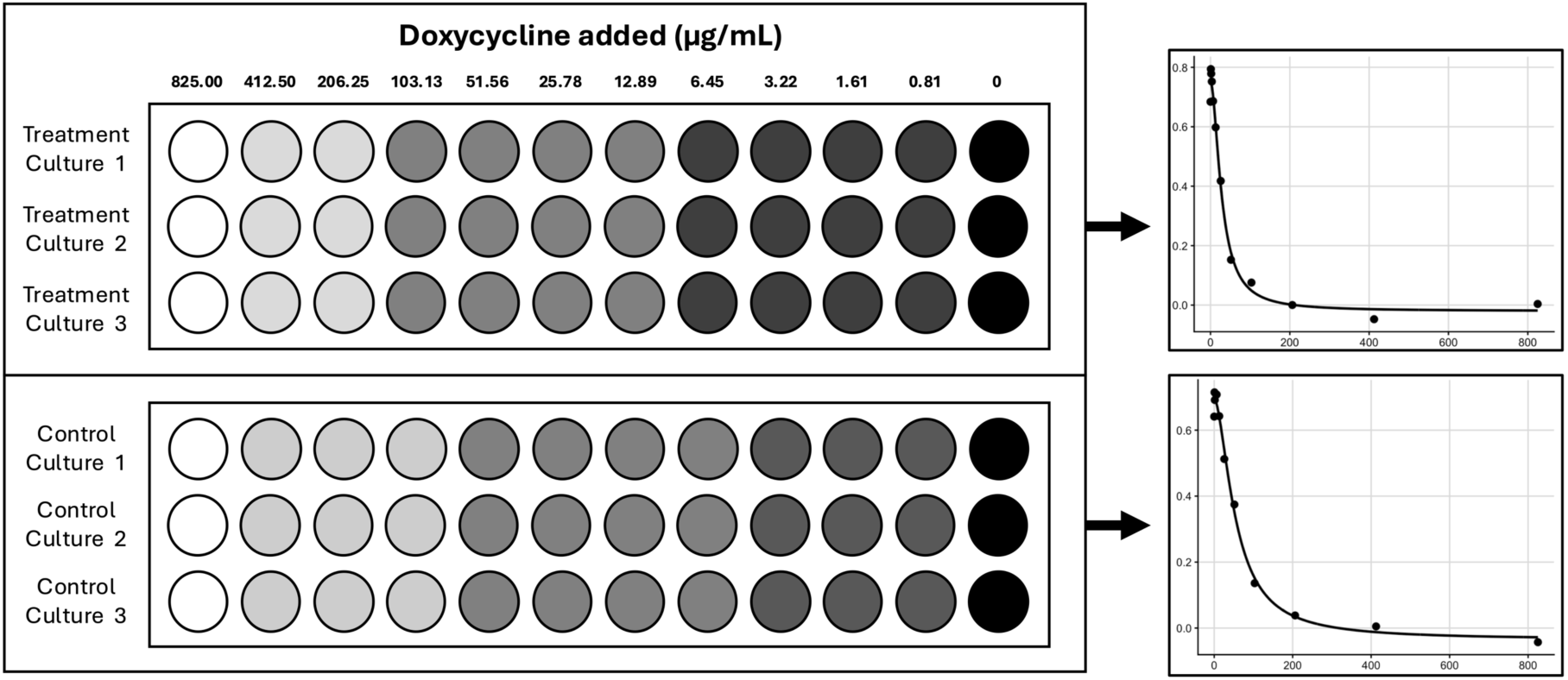
Fitness tests were used to assess changes in DOX resistance for *Lp14917* cultures and were analyzed visually and quantitatively. At regular intervals during the ALE, the fitness of DOX-exposed and control cultures was tested by incubating them in the presence of a sliding scale of DOX (wells 1-11). After incubation, growth was quantified by measuring the OD_600_, subtracting the initial absorbance value, and plotting net growth (the ΔOD_600_) as a percentage relative to an untreated control (well 12, no DOX) against DOX concentration. The ΔOD_600_ data was fit to a decreasing four-parameter Hill model, allowing determination of the inhibitory constant at any DOX concentration (such as IC_50_ values) and thus a complete inhibitory profile. Data for the control and DOX-exposed cultures from the ALE were treated and plotted independently.

Shortly after inoculation, the OD_600_ of the fitness test plates was measured and served as a starting point OD_600_ reading prior to incubation. Plates were incubated at 35°C in a stationary incubator for 24 hours, and the OD_600_ was measured again. The absorbance after 24 hours was subtracted from the absorbance before incubation. This difference represents the change in OD_600_ and thus growth over 24 hours. These differences were divided by the difference in OD_600_ of the untreated wells (no antibiotic, column 12) and multiplied by 100. This represents the percent change in OD_600_ in the DOX-treated wells relative to the change in OD_600_ of the untreated wells. Where growth is less in DOX-treated wells as compared to the untreated wells, this can be considered growth inhibition and an inhibitory concentration (IC) of DOX causing a certain percentage of inhibition can be roughly determined. For example, after 24 hours if the change in OD_600_ of well 4 is half that of well 12 (untreated), we can estimate that the IC_50_ for DOX of that specific colony is ∼103 µg/mL.

### Quantitative method for determining DOX resistance from fitness test data

While estimates of inhibition can be valuable and trends detected, we developed a method for quantifying DOX susceptibility from the fitness assays so as to compare resistance profiles for the two lineages (control vs. DOX-exposed). We constructed dose-response curves by plotting the 24-h growth range changes (subtracting the initial OD_600_, *y_min_* below, from the 24-hr OD_600_, *y_max_* below, to get an ΔOD_600_) as a function of antibiotic concentration, pooling measurements across the three replicates for each lineage. Control and DOX-exposed cultures were analyzed separately. For each generation and lineage, we fit the data using a decreasing four-parameter Hill Model,

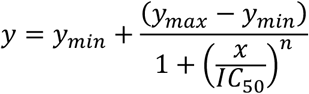

where *y_min_* and *y_max_* represent the lower and upper growth plateaus, *n* is the Hill coefficient, and *IC_50_* is the concentration producing a half-maximal response relative to the fitted plateaus (**Fig. 1**). Model parameters were estimated by nonlinear least squares, and goodness-of-fit was assessed using root mean square error (RMSE). The fitted values were then extracted for each generation and plotted against generation for the Control and DOX-exposed lineages on the same axis to visualize temporal changes in the susceptibility to DOX (**Fig. 2**). This framework enables direct tracking of resistance trajectories over time and facilitates comparison of the two lineages *via* their generation-specific estimates.

**Figure 2.**
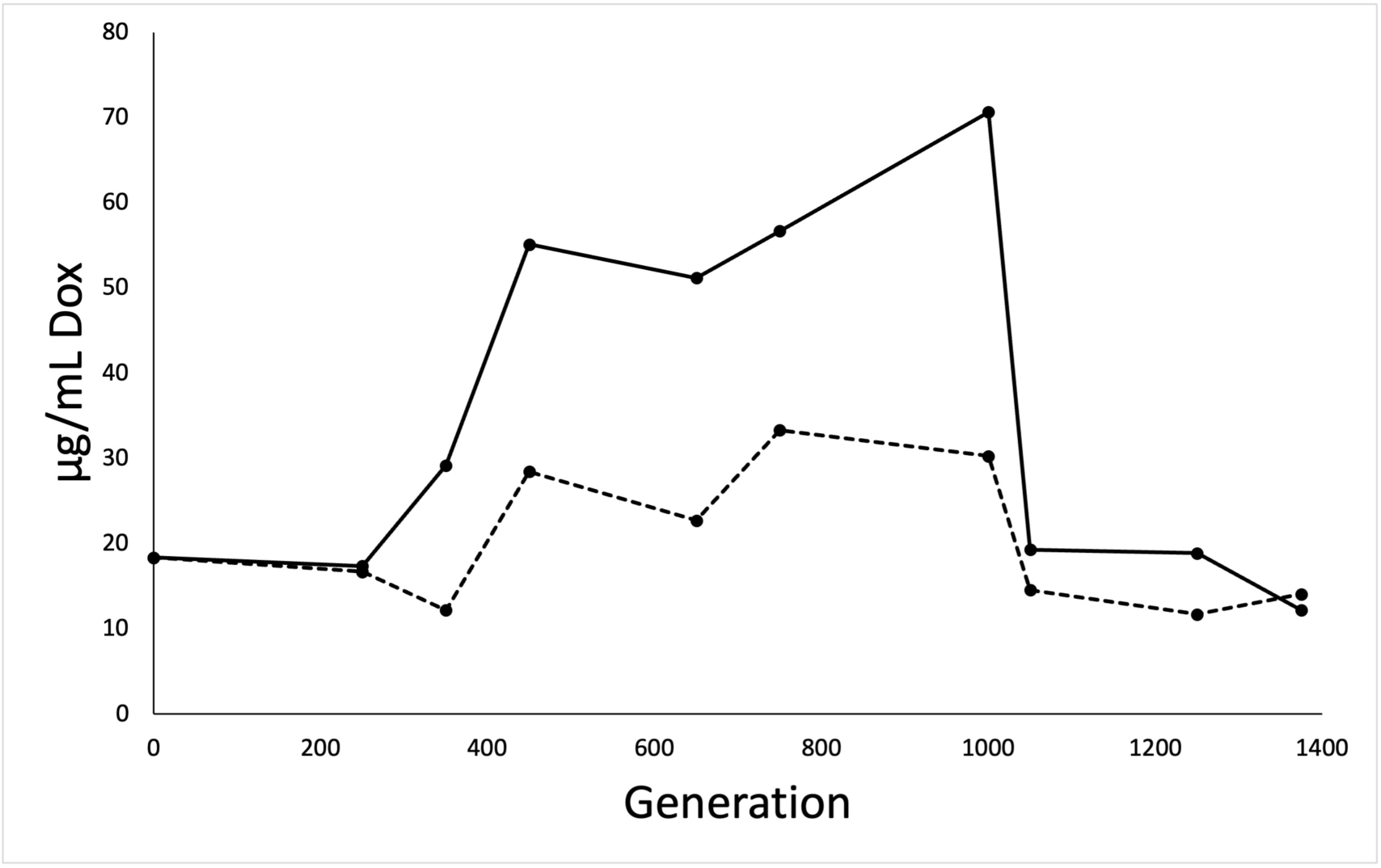
Upon sub-lethal exposure to DOX, *Lp14917* gains modest resistance to DOX and rapidly loses it after the selection pressure is removed. Cultures of *Lp14917* were propagated daily for nearly 1400 generations, and DOX fitness was regularly tested. Plotted here are the DOX IC_50_ values for control (dashed black line) and DOX-exposed (solid black line) cultures across the experiment. IC_50_ values were averaged from triplicate cultures and were calculated as shown in Fig. 1 and described in the text. At generation 1000, DOX was removed from the growth medium, and the experiment was continued to generation 1375. Fitness dramatically drops for the DOX-exposed cultures, suggesting loss of the acquired resistance.

### Whole genome sequencing of “archived” ALE generations

Genomic DNA was extracted from an overnight culture of *Lp14917* cells using the PureLink Genomic DNA Mini Kit (Invitrogen, ThermoFisher Scientific; Waltham, MA) with a slightly modified Gram Positive Bacterial Cell Lysate Protocol. Briefly, after cell lysis steps, we added 20 µL of RNase A and incubated at RT for 5 minutes. Additionally, prior to loading the homogenized lysate onto the purification columns, we clarified the samples by centrifugation (10 *k x g* for 10 minutes). Purified genomic DNA eluted from the column was quantified and checked for purity by UV spectrophotometry. As a quality control step, the variable 4 (V4) region of the 16s ribosomal subunit gene (16s rDNA) was amplified by PCR using gDNA samples of each of the timepoints as templates. Furthermore, Sanger sequencing of the V4 of the 16s rDNA from the Gen1000 extracted gDNA confirmed that, after the DOX exposure in the experiment, the strain was still identified as *L. plantarum* ATCC 14917.

Genomic DNA samples that passed our initial QC were shipped to Novogene (Sacramento, CA) for microbial whole genome sequencing (WGS). These were further QC-checked using a Qubit fluorometer (ThermoFisher Scientific; Waltham, MA) to determine concentration, and a 2100 Bioanalyzer System (Agilent Technologies; Santa Clara, CA) with high resolution agarose gel electrophoresis to determine purity and integrity. Only samples that passed all QC steps were selected for library construction and sequencing. These 19 samples represent controls and experimental populations across 5 time points (generations 0, 350, 750, 1000, and 1200).

### Library construction, quality control and sequencing

The genomic DNA was randomly sheared into shorter fragments. The obtained fragments were then end-repaired, A-tailed, and further ligated with Illumina adapters. The resulting fragments with adapters were size selected and PCR amplified before proceeding for purification. Purification was then conducted using the AMPure XP system (Beckman Coulter Life Sciences; Indianapolis, IN). The resulting library was assessed on the Fragment Analyzer System (Agilent Technologies; Santa Clara, CA) and quantified to 1.5 nM using a Qubit fluorometer as well as qPCR detection (ThermoFisher Scientific; Waltham, MA). The qualified libraries were pooled and sequenced on a NovaSeq6000 platform (Illumina; San Diego, CA) using the paired-end 150 base pair (PE150) method.

### Bioinformatic analysis of WGS data

The original fluorescence image data files obtained from the NovaSeq6000 were transformed to sequenced reads (raw reads) by base calling using CASAVA base recognition. Transformed raw data were stored in FASTQ format files (44, 45), which contain sequencing reads and corresponding base quality. The Phred score (Q_phred_= - 10log_10_(e)) defines the quality score of a base read and informs sequence quality distribution. A Phred score of 30 (Q30) corresponds to a correct read rate of 99.9%. All samples sequenced had 91-94% of reads fall within the Q30; the effective read rate was > 99% with an error rate of 0.03%. Sequence artifacts, including reads containing adapter contamination, low-quality nucleotides and unrecognizable nucleotides (reported as *N*), were identified and filtered by Fastp (version 0.23.1) (45). It was also used to perform basic statistics on the quality of the raw reads. Sequence reads were discarded in Fastp data processing if: (1) either paired read contains adapter contamination (> 10 nucleotides aligned to the adapter, allowing ≤ 10% mismatches); (2) more than 10% of bases are uncertain in either read; (3) if the proportion of low quality (Phred quality < 5) bases is over 50% in either read.

### Mapping reads to a reference sequence

The reference genome used for mapping reads is *L. plantarum* subspecies *plantarum* (Orla-Jensen) whole genome sequence (Genome ID= GCA_000143745.1) (46). This is the ATCC strain 14917 and is also known as Lp39. Valid sequencing data was mapped to the *L. plantarum* reference genome by Burrows Wheeler Aligner (BWA) (47) software to get the original mapping results stored in BAM format (parameter: mem - t 4 -k 32 -M). Then, the results were dislodged duplication by SAMtools(48) (parameter: rmdup) and Picard (http://broadinstitute.github.io/picard/). The mapping rate for all samples was between 98-99%.

### SNP/SNV variant detection

For each sample that underwent WGS, ∼1 GB of raw data was output. Based on the genome size of the *L. plantarum* reference genome (∼3.3 MB), sequence coverage would theoretically surpass 300X, which is quite sufficient coverage for single nucleotide polymorphism (SNP) or variant (SNV) detection. Across the 19 samples, the average sequence coverage depth was 350X, with a range from 244 to 539X. We detected the individual SNP variations using GATK (49) with the following parameter: ‘HaplotypeCaller--pair-hmm-gap-continuation-penalty 10 -ERC GVCF --genotyping-mode DISCOVERY -stand-call-conf 30’. To reduce the error rate in SNP detection, the SNP sets were filtered using the following criteria: QD < 2.0, FS > 60.0, MQ < 30.0, HaplotypeScore > 13.0, MappingQualityRankSum < −12.5, ReadPosRankSum < −8.0.

### Variant annotation

ANNOVAR(50) was used for functional annotation of variants. The UCSC database (http://genome.ucsc.edu) of known genes was used for gene and region annotations(51). The universal functional database used was the Cluster of Orthologous Groups (COG) protein database (52), and all predicted coding sequences were assigned to one of 25 functional groups (i.e. “carbohydrate transport and metabolism”). The basic steps for functional annotation included performing a BLAST search (53) between the predicted gene and various functional databases, and subsequently filtering the BLAST search results, selecting the hit with the highest overall score for each sequence.

### Verification of SNVs in the *rpsJ* gene

To verify identity and consequence to the amino acid sequence of the SNVs detected in the *rpsJ* gene from WGS, we coupled standard colony PCR amplification with Sanger sequencing. Frozen *Lp14917* from control and DOX-exposed cultures across generations 350, 750, and 1000 were quadrant streaked onto MRS agar plates (controls) or MRS agar plates supplemented with 3 µg/mL doxycycline (DOX-exposed). Plates were incubated at 35°C in a stationary incubator for up to 48 hours. Colonies (3 for controls, 5 for DOX-exposed) were selected for PCR, and a sterile tip was used to transfer a single colony into a PCR master mix (Phusion Green Hot Start II High-Fidelity PCR 2X Master Mix; ThermoFisher, Waltham, MA) in a 0.2 mL thin-walled tube. Forward and reverse primers to amplify the *rpsJ* gene were added to a final concentration in the PCR reaction of 2 µM. Nucleotide sequences for the primers were as follows: Forward, 5’-gcatggcaaagcaaaaaattcgtattc-3’, and Reverse, 5’-gcttaaagcttgatttcaatatctacacc-3’. PCR was performed on a SimpliAmp Thermal Cycler (Applied Biosystems, ThermoFisher, Waltham, MA), and cycling conditions were as follows: 3 min at 94°C for initial denaturation, 15 sec at 94°C for cycle denaturation, 15 sec at 58°C for primer annealing, 30 sec at 72°C for cycle extension, and 3 min at 72°C for final extension, with 30 total cycles. PCR reactions were analyzed for amplification of the *rpsJ* gene (amplicon size ∼306 bp) by horizontal agarose gel electrophoresis, composed of 1% w/v ultrapure agarose in 1 x TAE buffer and supplemented with 1:10,000 Sybr Safe DNA stain (Invitrogen ThermoFisher, Waltham, MA). Reactions positive for amplification were purified by silica column, washing, and centrifugation using the NucleoSpin Gel and PCR Cleanup Kit (Macherey-Nagel Inc., Allentown, PA). Purified PCR products (10 µL) were mixed with the *rpsJ* forward primer (5 µL) and shipped to Genewiz (Azenta Life Sciences, South Plainfield, NJ) for Sanger sequencing. Sequence text files and chromatograms were manually analyzed for the presence of SNVs previously observed from the WGS. Pairwise sequence alignment with EMBOSS Water (54) (http://ebi.ac.uk/jdispatcher/psa/emboss_water) was used to verify the presence of the expected SNV (or absence of an SNV if from a control colony) and to confirm other variants were not present. To compare sequence similarities in S10 proteins (the expression product of *rpsJ*) and to visualize variant positions on the S10 structure, a multiple sequence alignment was performed with CLUSTAL Omega (54) (version 1.2.4; https://www.ebi.ac.uk/jdispatcher/msa/clustalo) and a computed model provided by UniProt (55) (https://www.uniprot.org/uniprotkb/Q88XY7/entry) was analyzed.

### Data availability

Paired end reads from all the WGS data reported here are available at the NCBI BioSample Database (https://www.ncbi.nlm.nih.gov/biosample) in the Sequence Read Archive (SRA) (https://www.ncbi.nlm.nih.gov/sra), with the BioProject accession number, PRJNA1439351. The data can be accessed at https://www.ncbi.nlm.nih.gov/sra/PRJNA1439351.

## Results

### Adaptive Laboratory Evolution Experiment with *L. plantarum ATCC* 14917

Using disk diffusion, macro- and microbroth dilution assays, we established a minimal inhibitory concentration (MIC) for DOX for the inhibition of growth of *L. plantarum ATCC* 14917 (*Lp14917*). This value, 30 µg/mL, is in the range previously reported for *L. plantarum* strains isolated from fermented vegetables (22, 24). As we wanted to apply weak selection pressure to the bacteria, we exposed *Lp14917* to a sublethal concentration of DOX during an adaptive laboratory evolution (ALE) experiment. This entailed continuous culturing of *Lp14917* in MRS media with 1/10 MIC of DOX (so, 3 µg/mL). Six total cultures (3 DOX and 3 controls) were propagated daily. Based on our initial growth curves and previously reported ALE experiments with *L. plantarum*, we estimated 7 generations of *Lp14917* per day, thus 50 per week. As the experiment lasted roughly 20 weeks, we estimate ∼1000 total generations (42). Every ∼50 generations, aliquots of each culture were removed and frozen, and this represents a snapshot of the bacterial cells at that specific point in time. This archiving is common for ALEs as frozen cultures serve a “time machine” function for retrograde analysis(56).

### Determining and quantifying emergence of DOX resistance in *Lp14917* with fitness tests

Every 100-200 generations (once every 2-4 weeks), the “fitness” of both DOX-exposed and control cultures was determined by re-performing the microbroth dilution assay (**Fig. 1**). Instead of a MIC, here we focused on the IC_50_ value and to observe if it had shifted over the course of the ALE. In a 96 well plate, aliquots of *Lp14917* cells (either control or DOX-exposed) are mixed with DOX across a concentration range, and growth is measured as an absorbance (the OD_600_) at 0 and 24 hrs. For both controls and DOX-exposed cultures, the net growth (the ΔOD_600_) of each fitness test culture was expressed as a percentage of growth relative to an untreated culture control and graphically plotted against the concentration range of DOX used in the fitness test. In order to determine the exact point at which a precise reduction of growth occurred as an expression of DOX concentration, the absorbance data was fit using a decreasing four-parameter Hill model (**Fig. 1**).

From this, we established an IC_50_ per every generation tested (6 total) and directly compared IC_50_ values between DOX-exposed cultures to controls (**Fig. 2**, **Table 2**). The DOX IC_50_ for the DOX-exposed cultures increased to 29 µg/mL after 350 generations, which is similar to the MIC values for DOX we measured with disk diffusion and broth dilution assays. By the end of the ALE experiment (1000 generations), the IC_50_ had further increased to 70 µg/mL. This is a 3.9-fold increase from the beginning value (18 µg/mL) measured for the Gen0 original culture. Importantly, the IC_50_ values for control cultures essentially remain flat across the experiment, with an ending IC_50_ value of 30 µg/mL.

**Table 2.** DOX IC_50_ Values for Control and DOX-exposed *Lp14917* Across the ALE*.

| Generation<sup>†</sup> | Control | DOX-exposed |
| --- | --- | --- |
| 250 | 16.7 (2.3) | 17.4 (1.2) |
| 350 | 12.2 (1.7) | 29.2 (1.5) |
| 450 | 28.5 (1.8) | 55.1 (1.7) |
| 650 | 22.8 (1.4) | 51.2 (1.7) |
| 750 | 33.3 (1.8) | 56.7 (1.5) |
| 1000 | 30.3 (1.5) | 70.7 (1.6) |
| 1050 | 14.6 (2.4) | 19.3 (1.8) |
| 1250 | 11.7 (1.5) | 18.9 (1.5) |
| 1375 | 14.1 (2.4) | 12.2 (1.6) |

\*The IC<sub>50</sub> value is expressed as a concentration in µg/mL and rounded to the tenth. The number in parentheses is the root mean square error (RMSE) of the extracted values from the Hill model analysis.
<sup>†</sup>The original ALE starting culture had a DOX IC<sub>50</sub> of 18.4 µg/mL.

To test whether the increase in the IC_50_ value for DOX in the DOX-exposed cultures was reversible, we continued the ALE for 375 more generations, however, we omitted DOX from the media. Thus, both DOX-exposed and control cultures were grown and propagated the same (MRS broth alone). Again, we archived aliquots every 50 generations and performed a fitness test regularly. The IC_50_ value for DOX in the DOX-exposed cultures immediately decreased; after just 50 generations the IC_50_ values for DOX-exposed and control cultures were essentially the same (19 and 15 µg/mL, respectively) (**Fig. 2**, **Table 2**). Note that this value is nearly identical to the DOX IC_50_ value, 18 µg/mL, for the Gen0 original culture at the start of the ALE. The modest DOX resistance that had accumulated over 1000 generations was lost in less than a tenth of the time. Further fitness tests at Gen1250 and 1375 show the DOX IC_50_ values for DOX-exposed cultures to be steady and nearly identical to controls.

### Whole genome sequencing to identify variants that may be contributing to DOX resistance

To test whether the increase in the IC_50_ value for DOX in the DOX-exposed cultures has (at least in part) a genetic basis, we performed whole genome sequencing (WGS). Genomic DNA was extracted from archived *Lp14917* control and DOX-exposed samples from multiple colonies at 5 time points: Gen0 (the original ALE starting cultures), Gen350, Gen750, Gen1000, and Gen1200 (200 generations after DOX was removed). Variant analysis of the WGS data revealed 43 total exonic single nucleotide variants (SNVs) present in DOX-exposed colonies across the 1000 generations of the ALE experiment (**Table 3**); as a few of these are found at different time points, there are actually 34 *unique* SNVs. Importantly, with only one exception, these SNVs were not observed in either the Gen0 ALE starting cultures or the controls at any given time point. SNVs were located by identifying an alternate nucleotide call as compared to the reference nucleotide expected from a reference *Lp14917* genome sequence (Genome ID= GCA_000143745.1) (46). Having WGS data for multiple time points across the experiment, we can determine which and when certain SNVs arose (and, where applicable, when there is a reversion to the reference nucleotide). Sixteen of these emerge after Gen750, as they are only identified in sequences from Gen1000 colonies. Only 1 of these SNVs present at Gen1000 persisted in any Gen1200 colony. Surprisingly, the Gen1200 sequences revealed 24 unique SNVs that did not overlap variants identified in the ALE.

**Table 3.** SNVs Identified in DOX-exposed *Lp14917* Across the ALE.

| Variant/Position | Genotype* | Functional description | Class description |
| --- | --- | --- | --- |
| **Generation 350** |  |  |  |
| C -> T/67215 | 0/1 | Ribonucleotide reductase beta subunit | Nucleotide metabolism |
| C -> T/390012 | 0/1 | Ribosomal protein S10 rpsJ | Translation |
| A -> T/390016 | 0/1 | Ribosomal protein S10 rpsJ | Translation |
| C -> T/390349 | 0/1 | Transcriptional antiterminator | Transcription |
| **Generation 750** |  |  |  |
| C -> T/67215 | 1/1 | Ribonucleotide reductase beta subunit | Nucleotide metabolism |
| C -> T/390012 | 0/1 | Ribosomal protein S10 rpsJ | Translation |
| A -> T/390016 | 1/1 | Ribosomal protein S10 rpsJ | Translation |
| G -> T/390017 | 1/1 | Ribosomal protein S10 rpsJ | Translation |
| G -> A/390129 | 1/1 | Ribosomal protein S10 rpsJ | Translation |
| C -> T/390349 | 0/1 | Transcriptional antiterminator | Transcription |
| G -> A/197972 | 0/1 | Malonyl CoA-acyl carrier protein transacylase | Lipid metabolism |
| G -> A/198173 | 0/1 | Malonyl CoA-acyl carrier protein transacylase | Lipid metabolism |
| G -> A/541111 | 0/1;0/1;0/1<sup>†</sup> | LysM repeat | Cell wall biogenesis |
| C -> T/689750 | 0/1 | DedA/Tvp38 family (putative transporter) | Cell wall biogenesis |
| T -> A/55106 | 0/1 | EndoRNase, NYN family | Translation |
| G -> A/126954 | 1/1 | DNA-binding, GntR family | Transcription |
| G -> A/157876 | 0/1 | RNase adaptor RapZ | Signaling |
| G -> T/524611 | 0/1 | Carbamoylphosphate synthase, large subunit | Amino acid/nucleotide metabolism |
| G -> T/21523 | 0/1 | Glucosamine 6-phosphate synthetase | Cell wall biogenesis |
| C -> T/196230 | 0/1 | Hydroxymethylpyrimidine pyrophosphatase | Coenzyme metabolism |
| G -> A/36019 | 0/1 | Arginyl-tRNA synthetase | Translation |
| G -> A/104848 | 0/1 | *Homology to MFS transporter (?)* | ? |
| C -> A/187028 | 0/1 | *Homology to SpaA pilin-related protein (?)* | ? |
| C -> T/230743 | 0/1 | *Homology to GGDEF domain (?)* | ? |
| G -> A/314722 | 1/1 | *Homology to a hypothetical protein, WP_003640537.1 (?)* | ? |
| C -> A/313671 | 1/1 | *Homology to a hypothetical protein, WP_003641852.1 (?)* | ? |
| T -> G/118682 | 0/1 | *Homology to a hypothetical protein, WP_003640335.1 (?)* | ? |
| **Generation 1000** |  |  |  |
| C -> T/390012 | 1/1;1/1 | Ribosomal protein S10 rpsJ | Translation |
| G -> A/390129 | 1/1;0/1 | Ribosomal protein S10 rpsJ | Translation |
| G -> T/ 21523 | 1/1;0/1 | Glucosamine 6-phosphate synthetase | Cell wall biogenesis |
| G -> T/157876 | 1/1;1/1 | RNase adaptor RapZ | Signaling |
| C -> A/483670 | 1/1;0/1 | Asp-tRNAAsn/Glu-tRNA<sup>Gln</sup> amidotransferase | Translation |
| G -> T/192696 | 0/1;0/1 | GGDEF domain, Diguanylate cyclase | Signaling |
| G -> A/656201<sup>‡</sup> | 1/1;0/1 | Cell division ATPase FtsA | Cell Cycle Control |
| C -> T/197765 | 0/1 | Malonyl CoA-acyl carrier protein transacylase | Lipid metabolism |
| G -> A/276190 | 0/1 | Lysozyme M1, GH25 family | Cell wall biogenesis |
| C -> T/ 388028 | 0/1 | Phosphohistidine swiveling domain of PEP-utilizing enzymes | Signaling |
| G -> T/537669 | 0/1 | SOS-response transcriptional repressor LexA | Signaling |
| C -> A/435174 | 0/1 | Malate/lactate dehydrogenase | Energy conversion |
| T -> G/161050 | 0/1 | ECF-type transporter transmembrane protein EcfT | Coenzyme transport |
| G -> T/165985 | 0/1 | Beta-glucosidase/Beta-galactosidase | Carbohydrate metabolism |
| C -> A/286503<sup>^</sup> | 0/1 | *Homology to phage portal protein (?)* | ? |
| G -> T/43917 | 0/1 | *Homology to aryl-sulfate sulfotransferase (?)* | ? |

\*0/1 = heterozygous, where a mixture of reference and variant alleles were called; 1/1 = homozygous for the variant allele (>95% of calls were for the variant)
<sup>†</sup>Where multiple genotypes are listed, this indicates the variant was identified across multiple cultures (colonies)
<sup>‡</sup>A G -> T variant was identified at this location in one of the control colonies, with a 0/1 genotype
<sup>^</sup>This is the only variant that arises in the ALE (sometime prior to generation 1000) that persists to generation 1200 (cultures from generation 1000 to 1200 were grown in the absence of doxycycline)

### Annotation of the variants identified from sequencing

Functional annotation of the SNVs, mainly using the Cluster of Orthologous Groups (COG) database(52), shows that the highest frequency of variation occurs in genes associated with either transcription or translation (**Table 3**). Of the 34 unique DOX-associated SNVs, 4 were identified in the *rpsJ* gene, which encodes the ribosomal protein, S10 (**Table 4**). Previous studies have shown that S10 interacts with a specific helix within the 16s rRNA subunit (i.e. h31 in *Klebsiella pneumoniae* and *Escherichia coli*), primarily mediated by a flexible loop composed of amino acids 50-60 (29, 33, 35) (**Fig. 3**). This interface lies adjacent to (though not overlapping with) the tetracycline class antibiotic binding site, near ribosomal translation site A. The *rpsJ* SNVs identified in this study from WGS were H56Y, K57M, K57I, and S94N (**Table 4**), the first 3 located in the aforementioned flexible loop that contacts the 16s subunit. The S10 H56Y SNV is observed at Gen350 and persists at Gen750 and 1000 whereas the K57I variant is first observed at Gen750 but is absent at Gen1000. We assume this is a reversion event back to the reference nucleotide at this position. Although the K57I variant is not observed at Gen1000, two other variants are observed, H56Y and S94N.

**Figure 3.**
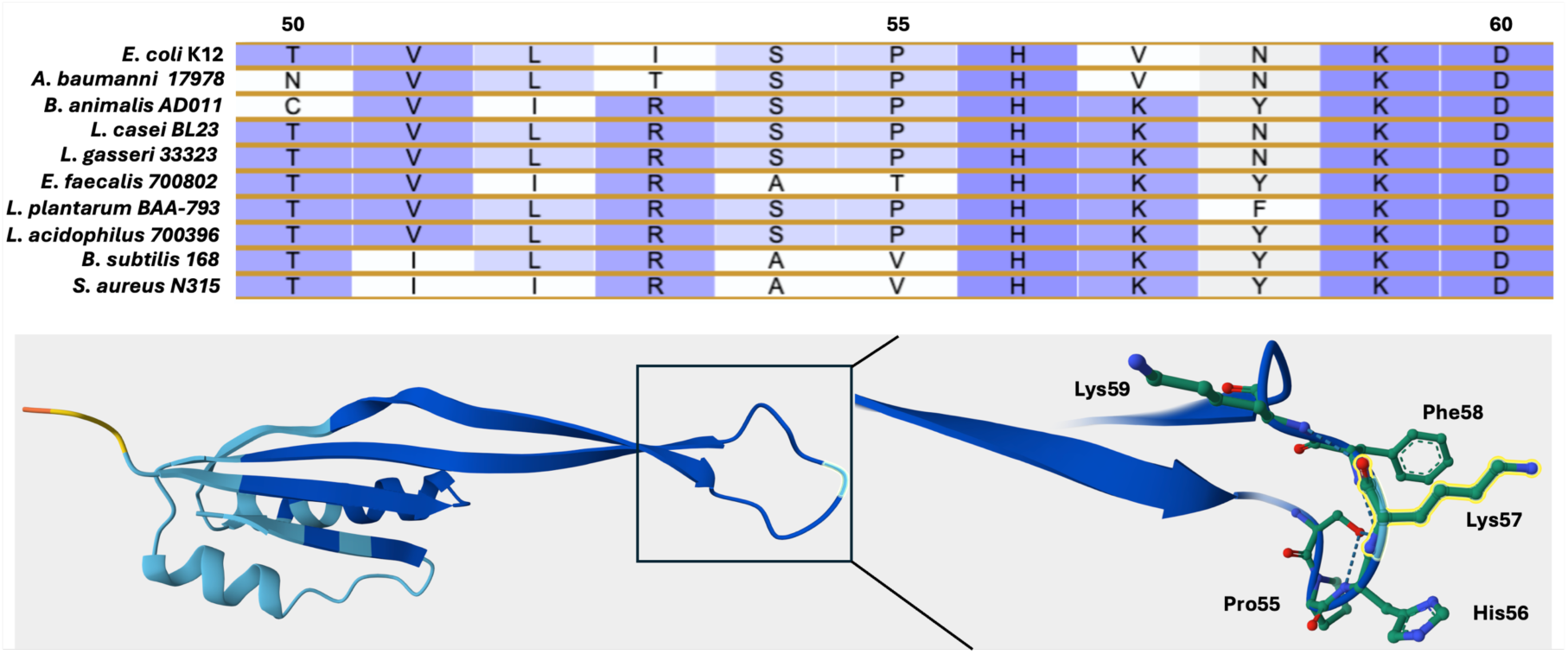
Nearly all *rpsJ* variants map to the tip of a large, unstructured loop between strands B and C of the S10 protein. Amino acid sequence alignment for S10 proteins from Gram negative (*E. coli* and *A. baumanni*) and Gram positive (all others) species shows general conservation in this B-C loop, with identical residue for all at positions 56, 59, and 60 (*E. coli* numbering). Note the dichotomy for residue 57: V for Gram negatives, K for Gram positives. This is one of the most common mutation sites in the rpsJ gene associated with tetracycline class resistance. Multiple sequence alignment was performed using CLUSTAL Omega, version 1.2.4 (https://www.ebi.ac.uk/jdispatcher/msa/clustalo). To visualize the S10 protein structure from *L. plantarum BAA-793* (a strain closely related to *Lp14917*), an AlphaFold2 computed model was used (AF-Q88XY7-F1) as provided by the UniProt (https://www.uniprot.org/uniprotkb/Q88XY7/entry). Model is rendered in cartoon format, and the backbone is colored using a per-residue confidence score, from dark blue (very high confidence) to orange (very low confidence). Only the side chains for residues 55-59 are shown, and they are rendered as ball-and-stick model. Side chain atoms are colored by element: carbon (green), nitrogen (blue), and oxygen (red). Potential intramolecular hydrogen bonding is shown as dashed lines.

**Table 4.**
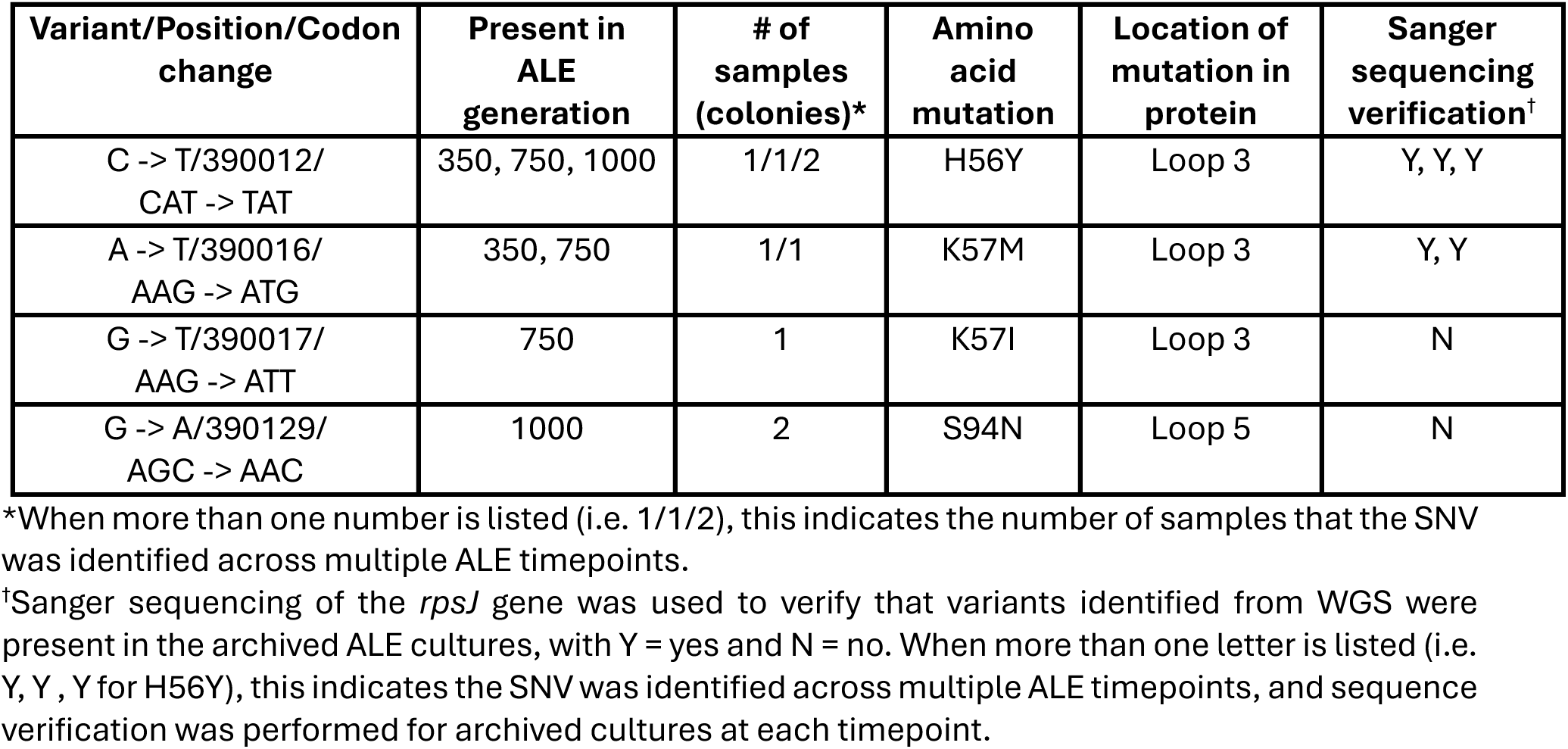
SNVs Identified from the WGS for *rpsJ*, which encodes ribosomal protein S10.

### Verification of rpsJ variants by colony PCR and sequencing

Of the 4 SNVs in *rpsJ* that were observed from the WGS, 3 variants (H56Y, K57M, and K57I) could be verified by colony PCR and Sanger sequencing of the amplified *rpsJ* gene (**Tables 4** and **5**). One of the variants, H56Y, was identified in multiple colonies from archived cultures from Generations 350 and 1000. However, this variant was not observed in Gen750 colony PCR sequences, though it was identified in the WGS. Furthermore, the S94N variant was also not observed in Gen1000 colony PCR sequences, yet it had been identified in the WGS. In contrast, 3 variants, all at codon position 57 (K57M, K57N, and K57T), could be observed in colony PCR sequences, however, these had not been identified by WGS (**Table 5**). In the case of K57M, this variant was observed in WGS for Gen350 and Gen750, but not in Gen1000; from the colony PCR, it could be observed in Gen1000.

**Table 5.**
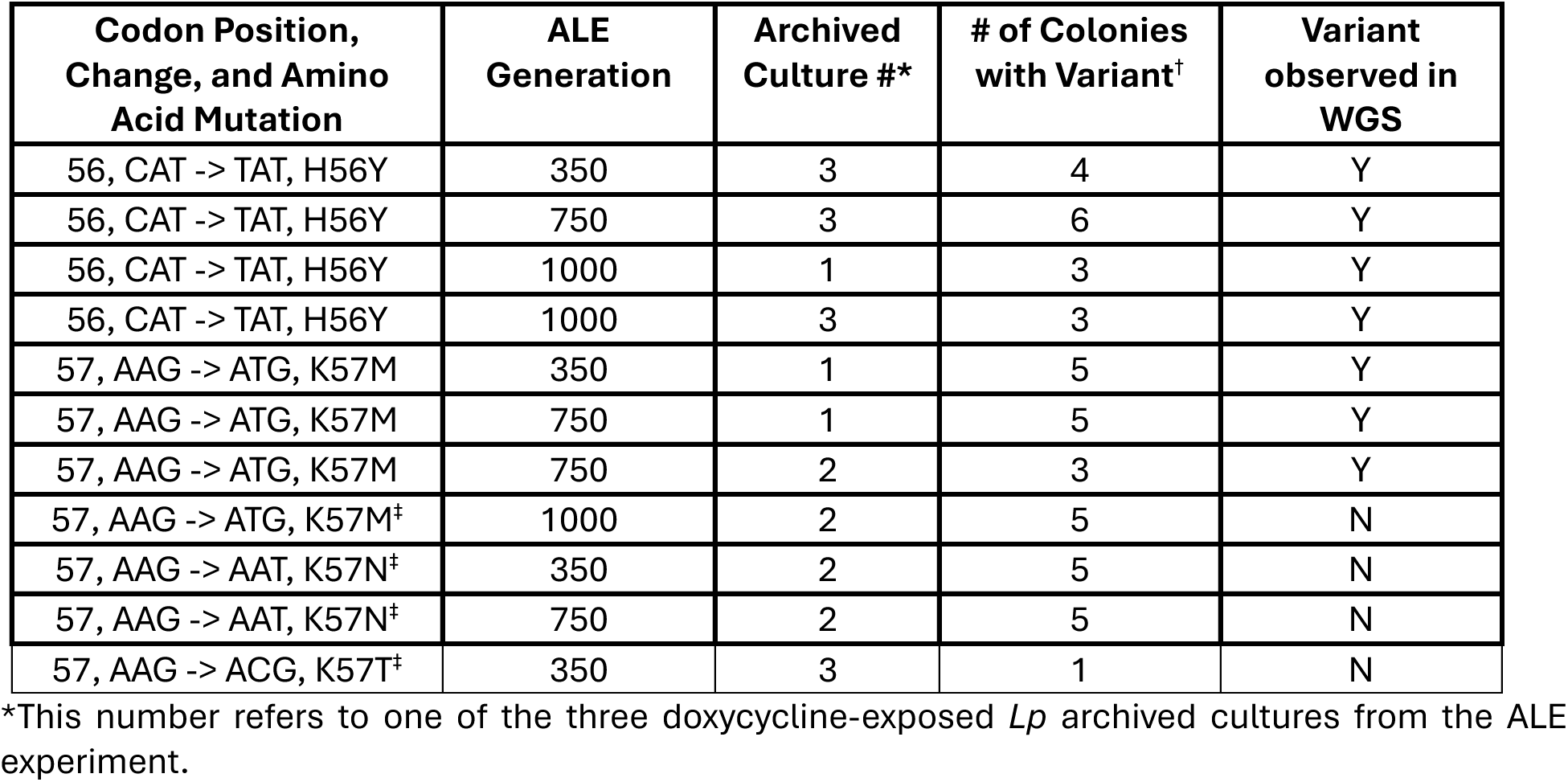
SNVs Identified at Positions 56-57 of *rpsJ* using Sanger Sequencing.

| Codon Position, Change, and Amino Acid Mutation | ALE Generation | Archived Culture #* | # of Colonies with Variant<sup>†</sup> | Variant observed in WGS |
| --- | --- | --- | --- | --- |
| 56, CAT -> TAT, H56Y | 350 | 3 | 4 | Y |
| 56, CAT -> TAT, H56Y | 750 | 3 | 6 | Y |
| 56, CAT -> TAT, H56Y | 1000 | 1 | 3 | Y |
| 56, CAT -> TAT, H56Y | 1000 | 3 | 3 | Y |
| 57, AAG -> ATG, K57M | 350 | 1 | 5 | Y |
| 57, AAG -> ATG, K57M | 750 | 1 | 5 | Y |
| 57, AAG -> ATG, K57M | 750 | 2 | 3 | Y |
| 57, AAG -> ATG, K57M<sup>‡</sup> | 1000 | 2 | 5 | N |
| 57, AAG -> AAT, K57N<sup>‡</sup> | 350 | 2 | 5 | N |
| 57, AAG -> AAT, K57N<sup>‡</sup> | 750 | 2 | 5 | N |
| 57, AAG -> ACG, K57T<sup>^</sup> | 350 | 3 | 1 | N |

\*This number refers to one of the three doxycycline-exposed *Lp* archived cultures from the ALE experiment.
<sup>†</sup>This number refers to the quantity of *Lp* colonies from the plated archived cultures that had a variant at position 56 or 57 in the *rpsJ* gene, as determined by Sanger sequencing. For most of the entries listed, up to 5 colonies were analyzed.
<sup>‡</sup>These variants were not observed in the original archived cultures sequenced by WGS. Note that a K57M variant was identified by Sanger sequencing in generation 1000. Even though this variant was observed in generations 350 and 750 from the WGS, it was not observed at generation 1000.

No *rpsJ* variants were detected in sequences (either by WGS or Sanger) from the original cultures (Gen0) or in same-generation controls. Additionally, variants at positions 57 and 94 could not be detected in any of the Gen1200 sequences. However, one of the DOX-exposed cultures retained the H56Y mutation at Gen1200. This wasn’t identified in the WGS but was detected in the Sanger sequencing. The other Gen1200 DOX-exposed cultures were WT at this position.

### Other variants identified at multiple time points

Twenty-eight additional SNVs were identified from the WGS across generations 350, 750, and 1000 (**Table 3**). Only 4 persisted in multiple generations: ribonucleotide reductase (RNR), BglG family transcriptional antiterminator, LysM repeat, and glucoside-6-phosphate synthase. The first is critical in the maintenance of NTP/dNTP pools while the second has a role in transcription regulation, allowing RNA polymerase to process beyond an initial termination sequence in a gene being transcribed. The latter two (LysM and Gln-6-P synthase) both localize to bacterial cell walls and assist in degradation (LysM) or synthesis of (Gln-6-P synthase) peptidoglycan matrix. Eight SNVs could not be sufficiently annotated; manual searches in NCBI using a gene ID have allowed putative assignment, sometimes only to a hypothetical protein. Furthermore, a single insertion-deletion (indel) not found in controls could be identified: a 69-nucleotide insertion in an exonic region, within the gene encoding tRNA-Arg.

## Discussion

Through an ALE-based experiment using weak selective pressure, we exposed the important probiotic *L. plantarum* (strain ATCC 14917) to sublethal concentrations (1/10 MIC) of the tetracycline antibiotic, DOX. Continual culturing was done over 1000 generations (∼5 months), with propagations every 24 hours into fresh media. To measure growth and determine a change in antibiotic resistance, we performed regular fitness tests. Though visual interpretation of growth is standard to determine MIC values, we complemented this with quantitative measures of growth using a plate reader spectrophotometer and recording optical density (OD_600_) of the bacterial cultures. Growth could be plotted against DOX concentration, and a Hill model curve fitted, quantifying how much growth has occurred at a given DOX concentration. A complete inhibitory profile for a strain versus a specific antibiotic can be precisely estimated by this method. We used the IC_50_ value to inform our interpretation of the ALE and fitness test results.

Within ∼6 weeks, the DOX-exposed cultures of *Lp14917* could survive higher concentrations of the antibiotic compared to the initial ALE start culture (Gen0) and to same generation controls. By the end of the experiment, we observed a ∼4-fold relative increase in the DOX IC_50_ for the Gen1000 DOX-exposed cultures. Though modest, it suggests sublethal concentrations can provide selection pressure and drive antibiotic resistance. Upon removal of the antibiotic, we observed a loss of resistance within ∼50 generations, a much faster rate in which it was acquired, showing resistance can be reversible and rapid. One potential basis for this acquired resistance could be at the gene/genomic level. Whole genome sequencing (WGS) and subsequent analysis of the reconstructed and annotated genome revealed 43 total exonic single nucleotide variants (SNVs) and 1 insertion/deletion (indel) unique to DOX-exposed cultures across the ALE. These variants were not identified in the original culture (Gen0), same generation controls, or nearly any of the Gen1200 cultures, 200 generations after the antibiotic had been removed. Several SNVs were mapped to the *rpsJ* gene, which encodes a ribosomal protein (S10) that assists with elongation during translation. Most of these could be verified by colony PCR and Sanger sequencing, with an additional 4 SNVs being identified this way. All the *rpsJ* variants are non-synonymous, causing an amino acid change in the translated polypeptide sequence. Even though DOX resistance could have multiple causes, it is at least in part correlated to one or more of the identified *rpsJ* variants. For example, compared to the original culture at Gen0 and same generation controls, we observe increased DOX IC_50_ values for DOX-exposed cultures at the generations for which we have WGS data (Gen350, 750, and 1000) (**Table 2** and **Fig. 2**), and there are *rpsJ* variants identified at each of these time points. However, the root cause for DOX resistance may rely on other factors, including but not limited to variants in other genes and/or changes in gene expression. The implication of *rpsJ* variants on tetracycline-class antibiotic resistance, including this study, is discussed in detail below.

As far as we know, this is the first ALE experiment with *L. plantarum* using sublethal tetracycline-class antibiotic as the selection pressure. There are several previous ALE studies of *L. plantarum* using sublethal antibiotics of other classes, including the beta-lactam ampicillin(57), and the aminoglycosides streptomycin (38, 39) and gentamicin (40, 41) (**Table 1**). All the aminoglycoside experiments were performed over 25-30 days, with the concentration of antibiotic in ALE cultures being adjusted based on fitness tests. For example, in the gentamicin experiment detailed in Dong *et al.* (2019) (41), an IC_50_ for gentamicin was determined in their initial antibiotic susceptibility tests, and this value was used as the starting gentamicin concentration in the ALE experiment. This concentration was increased 1.5-fold every 5 days, as fitness tests determined the “new” MIC and informed to what extent the gentamicin concentration should be adjusted^7^. For two of the aminoglycoside experiments, one with streptomycin^5^ and one with gentamicin^7^, once a stable new MIC had been reached, the antibiotic was removed from the ALE cultures, and it was observed that the MIC decreased significantly and rapidly but not back to its original level, showing at least partial reversibility to the acquisition of resistance. The ampicillin ALE experiment continued for much longer (12 months, ∼2400 generations), and the antibiotic concentration across the entire experiment was fixed at 0.5 of the MIC (57). We blended the format of all these studies as a model for our ALE here with doxycycline. Like Cao *et al.* (2020) (57), we ran the experiment at a fixed antibiotic concentration and the duration lasted several months; ultimately, we stopped the experiment at nearly 1400 generations. Similarly to Zhang *et al.* (2018) (39) and Dong *et al.* (2019) (41), we removed antibiotic from the ALE cultures and observed a rapid decrease in antibiotic fitness, suggesting a reversible loss of the resistance acquired in the experiment.

Previous ALEs where resistance to a tetracycline class antibiotic was selected for has been demonstrated in a number of bacterial species, including Gram positives such as *Staphylococcus aureus* (32) and *Streptococcus pneumoniae* (34) (both to tigecycline, or TGC) as well as Gram negatives such as *Escherichia coli* (29) (to tetracycline, or TET, and TGC), *Vibrio cholerae* (37), and *Neisseria gonorrhoeae* (36) (both to DOX). Most tetracycline-class resistance mechanisms are based on ribosomal protection (such as the TetM elongation factor protein in *E. coli*) or several families of efflux pumps, such as TetL or AcrAB-TolC (24, 28, 29). However, resistant mutations have also been mapped to genes encoding 16s rRNA and associated ribosomal proteins, such as in *rpsC* and *rpsJ,* which encode the S3 and S10 proteins, respectively. This has been especially the case for ALEs where TGC resistance has been cultivated (29) and even in clinical isolates where TGC resistance has been observed (30). Both the S3 and S10 proteins interact with helices in the 16s rRNA but don’t directly contact the A site or bind TGC. All reported S10 mutations reside in a long and unstructured loop (amino acids 50-60) adjacent to the TGC binding site and in contact with the h31 helix in the 16s rRNA. Indeed, to identify different rpsJ mutations possible, Beabout *et al.* (2015) performed a TGC-based ALE with several strains across 4 bacterial species (*Acinetobacter baumanni, Enterococcus faecium, E. coli,* and *S. aureus*). About 75% of the adapted and tested populations had at least one mutation in *rpsJ*, and all of them were observed in the aforementioned loop, with the highest concentration being at position 57 (33).

This agrees with the *rpsJ* variants defined in this study, identified by either WGS or colony PCR paired with targeted Sanger sequencing. Of the 13 total *rpsJ* variants, all are at positions 56-57 except 1 instance of a S94N variant in two of the Gen1000 DOX-exposed cultures (**Tables 4** and **5**). As this variant could not be verified by follow-up PCR and sequencing and has not been described previously, we lack confidence on how much importance to assign it. All the other variants map onto the B-C loop of S10, between strands B and C. This region possesses fairly high sequence similarity between residues 50-60, even when comparing Gram positive and Gram negative species (**Fig. 3**) (54, 55). An exception is position 57, where lysine is preferred in Gram positives and valine is found in Gram negatives. Upon DOX exposure, the S10 for *L. plantarum* was most often found with a K57M mutation, and this was seen across all generations of the DOX-exposure phase of the ALE (Gen350, 750, and 1000). This is one the most common S10 mutations associated with tetracycline-class antibiotic resistance in Gram positives (33). However, other variants have been identified at this position, and we observed K57I and K57N mutations from our sequencing data. Substituting lysine for any of the variant alleles observed here (M, I, N, or T) would neutralize a positive charge, and this has been implicated to be important for maintaining an electrostatic interaction with rRNA helices h31 and 34. This may in turn help to stabilize antibiotic binding near the A site (28); thus, losing this K57:rRNA helix charge-charge interaction may weaken antibiotic binding. Of the variants at position 57 reported here, all reduce the van der Waals (vdW) volume at this position (K is 135Å^3^, whereas the others are 124 (M), 124 (I), 96 (N), and 93Å^3^ (T), respectively). Perhaps volume reduction paired with charge neutralization causes the helices to reposition and interfere with DOX binding. Interestingly, by chromosomal integration of *rpsJ* mutant alleles into *E. coli*, it was found that exchanging V57 (the preferred amino acid in Gram negatives) to lysine (preferred in Gram positives) made the *E. coli* more susceptible to tetracycline (29). An *rpsJ* mutant identified in *Enterococcus faecalis*, known as S613-S10^R53Q-τι54-57ATHK^, has a missense substitution at position 53 and a 12-nucleotide deletion that doesn’t cause a frameshift. This mutant was shown to grow to a similar density in TGC-free media, as compared to the ancestral susceptible strain. Comparable growth rates suggests that, at least this specific mutation, likely presents a low fitness cost and could persist as a stable allele in a population (33). The growth rate study in the absence of TGC was performed over 12 hours (∼2-3 generations) and is of course likely not enough time to determine whether the mutation would actually be suppressed and the predominant allele in the population would revert to wild type in the absence of the selection pressure. From the WGS reported here, none of the *rpsJ* variants at K57 persist at Gen1200, ∼200 generations after the removal of DOX from the experimental cultures. Fitness tests at Gen1050, after 1 week of DOX-free growth, show a significant drop in the IC_50_, suggesting a very rapid loss of DOX resistance.

Like the K57M mutation, we observed numerous instances of an H56Y mutation across Gen350, 750, and 1000. Unlike K57M, there is little previously documented about this specific mutation; to our knowledge, it has only been reported once, found in clinical isolates of *E. faecalis* with omadacycline resistance (58). A histidine at this position is well conserved across bacteria. As for its role in DOX resistance, the vdW volume of tyrosine (141Å^3^) is significantly larger than that of histidine (118Å^3^), so perhaps the added bulkiness of the aromatic hydroxyl-substituted ring sterically conflicts with DOX at its ribosomal binding site. Surprisingly, targeted Sanger sequencing for *rpsJ* from one of the DOX-exposed cultures at Gen1200 (200 generations after DOX had been removed from the growth media) revealed an H56Y mutation. This mutation was not observed in the WGS for any of the cultures at Gen1200, and the two other DOX-exposed cultures at Gen1200 did not have this mutation. As none of the DOX-exposed cultures at Gen1200 have mutations persist at position 57, this suggests a higher fitness cost for variation at K57 than at H56. As both of these variants are found in Gen1000 cultures, where the highest DOX IC_50_ was measured, perhaps there is a synergistic effect on resistance underscored by mutation at these adjacent positions.

Concern about antibiotic resistance is typically nominal for probiotics, and much of this is because probiotics have rarely been observed to develop into a pathogen and become harmful to their hosts. Lactobacilli are classified as Generally Regarded As Safe (GRAS) microbes by the US FDA (59). However, in persons with certain comorbidities or previous pathological conditions, there are reports of clinical isolates of Lactobacilli associated with medical problems and diseases, often due to virulence factors. This can be concerning, not just for the infecting Lactobacilli, but also as they could harbor and be a reservoir for ARGs and virulence factors that could be horizontally transferred to more classical pathogens (22). On the other hand, when probiotics do evolve antibiotic resistance, the predominant variants and mutants that arise are sometimes not within or near mobile genomic elements or on plasmids that could be horizontally transferred (24).

Could probiotic strains with enhanced resistance be advantageous for synergistic probiotic:antibiotic treatments of certain gastrointestinal diseases? Antibiotics are widely known to exacerbate GI distress, leading to diarrhea and discomfort, along with gut dysbiosis as the antibiotic may non-specifically target beneficial members of the gut microbiome. A fairly recent trend has been to pair antibiotic therapy with ingestion of probiotics, especially blends of several LAB species, to maintain gut microbiome diversity and enhance Firmicutes/Bacteriodetes phyla over Enterobacteia ones (60). The former group is associated with better GI health outcomes while the latter group represent many of the important enteric pathogens and are known harbors of ARGs. Indeed, long-term consequences of antibiotic therapy can even lead to persistence of ARGs (61). A recent placebo-controlled study showed that when adults pair oral antibiotic treatment with a cocktail of 4 probiotic species and 1 yeast commensal, their normal GI microbial diversity is preserved and the abundance of ARGs in the gut is significantly reduced (62). A concern is the antibiotic targeting the probiotic species themselves, reducing their numbers and discounting their effectiveness. One solution is to pair antibiotics with a probiotic that is not susceptible to it, however, this can be impractical. Another solution is to “protect” the probiotic from the antibiotic until it is within the gut and can be established; for example, probiotics encapsulated in an alginate “cage” shields them from tobramycin yet allows by-products and secretions from the probiotic (known as postbiotics) to diffuse outward (63). Perhaps an additional solution to ensure probiotic survival is to pair probiotics with acquired resistance to the antibiotic used for treatment. In our present ALE study, we observed *Lp14917* attain ∼4-fold increase in the IC_50_ for doxycycline. Though modest, it does result in an adapted strain with enhanced resistance to a specific antibiotic.

While the *rpsJ* variants identified and described here likely correlate to some of the DOX resistance observed, there could be many contributions to enhanced resistance, including SNVs in other genes, indels and structural variations in the genome, transcription rate changes in *rpsJ* and other DOX-associated genes, and protein stability effects, such as those described by Cao *et al.* for *L. plantarum* exposed to ampicillin (57). A future goal is to use RT-qPCR with the archived *Lp14917* cells from the ALE (our “fossil record”) to monitor transcription level changes to *rpsJ*, *tet* resistance genes, and a few of the other SNV targets identified by the WGS, such as RNR and the BglG transcriptional antiterminators. Additionally, chromosomal integration of the *rpsJ* variants identified here could provide a powerful method by which to test *in vivo* the direct effect on DOX resistance of any single mutation in this gene, and it would be expressed from the background of an otherwise WT strain with no other gene variants. This targeted replacement of *rpsJ* has been successfully employed in a number of Gram positive and Gram negative species (29, 33).

## Acknowledgments

BP and BCB are grateful for funding through the Arts and Sciences Program for Independent Research Experiences (ASPIRE) at Samford University. We thank several undergraduate research students who helped at various points in this project, including Ansley Applestone, Barron Cate, Ben Chandler, Nicholas Hammond, and Camryn Pierce.

